# A monocarboxylate transporter rescues frontotemporal dementia and Alzheimer’s disease models

**DOI:** 10.1101/2023.06.25.546435

**Authors:** Dongwei Xu, Alec Vincent, Andrés González-Gutiérrez, Benjamin Aleyakpo, Sharifah Anoar, Ashling Giblin, Magda L Atilano, Mirjam Adams, Dunxin Shen, Annora Thoeng, Elli Tsintzas, Marie Maeland, Adrian M Isaacs, Jimena Sierralta, Teresa Niccoli

## Abstract

Brains are highly metabolically active organs, consuming 20% of an organisms’ energy at resting state. A decline in glucose metabolism is a common feature across a number of neurodegenerative diseases. Another common feature is the progressive accumulation of insoluble protein deposits, it’s unclear if the two are linked.

Glucose metabolism in the brain is highly coupled between neurons and glia, with glucose taken up by glia and metabolised to lactate, which is then shuttled via transporters to neurons, where it is converted back to pyruvate and fed into the TCA cycle for ATP production. Monocarboxylates are also involved in signalling, and play broad ranging roles in brain homeostasis and metabolic reprogramming. However, the role of monocarboxylates in dementia has not been tested.

Here, we find that increasing pyruvate import in *Drosophila* neurons by over-expression of the transporter *bumpel*, leads to a rescue of lifespan and behavioural phenotypes in fly models of both frontotemporal dementia and Alzheimer’s disease. The rescue is linked to a clearance of late stage autolysosomes, leading to degradation of toxic peptides associated with disease. We propose upregulation of pyruvate import into neurons as potentially a broad-scope therapeutic approach to increase neuronal autophagy, which could be beneficial for multiple dementias.

## Introduction

The brain is a highly metabolically active organ, using about 20% of the energy consumed by the human body [1]. In neurons glycogen storage is very reduced or non-existent, therefore they rely on a continuous supply of glucose to fuel their high energy requirements. Most of this glucose is used to maintain synaptic activity [2]. Interestingly, dementia progression is associated with a drop in glucose metabolism in the brain. This is well established in Alzheimer’s disease (AD), where glucose hypometabolism precedes the onset of clinical symptoms of disease [3], worsens with disease progression and mirrors the pattern of brain atrophy [4]. A reduction in expression of several glucose transporters has been observed in the brains of mouse AD models [5] and of patients [6]. Alzheimer’s disease is caused by the accumulation of extracellular plaques composed of Amyloidß (Aß) peptides, derived from the mis-processing of the Amyloid Precursor Protein (APP). Aß 1-42 is a major component of plaques [7], and dominant genetic mutations linked to familial AD, lead to an increase in its production [8]. Increase amounts of Aß 1-42 in animal models lead to pathology and neurodegeneration [9], underlying its causal role in disease development.

Patients with frontotemporal dementia (FTD), the second most common early onset dementia, also present with glucose hypometabolism in the frontal and temporal areas [10, 11]. This hypometabolism pre-dates dementia in FTD linked Granulin (GRN) mutant carriers [12], spreads as disease progresses [13] and in Microtubule-Associated Protein Tau (MAPT) -linked FTD patients closely maps to areas of toxic MAPT accumulation [14], suggesting it closely tracks disease development. One study also noted that glucose hypometabolism seemed to precede structural abnormalities [15]. Moreover, carriers of a MAPT haplotype linked to an earlier onset show reduced glucose metabolism in frontal areas [16] and diabetes is an independent risk factor in FTD development [17], suggesting that glucose metabolism might drive disease development. Very recently, an FTD mouse knock-in model carrying a Triggering Receptor Expressed On Myeloid Cells (TREM2) mutation, showed a significant reduction in glucose metabolism, providing further support to the link between FTD and glucose metabolism [18]. A hexanucleotide G4C2 repeat expansion located in the first intron of the *C9orf72* gene is the most common genetic cause of both amyotrophic lateral sclerosis (ALS) and FTD [14]. C9 repeats are transcribed both in the sense and antisense direction to generate highly stable, repetitive mRNAs which are translated in all open reading frames through a non canonical, repeat associated non-ATG mediated form of translation (RAN translation), to generate 5 different dipeptide repeat proteins (DPRs): Glycine-Alanine (GA), Glycine Proline (GP), Glycine Arginine (GR), Proline Alanine (PA), Proline Arginine (PR) [19]. The arginine containing proteins are the most toxic in the majority of model organisms where a head to head comparison has been described [20]. Patients with C9 mutations, like other ALS and FTD patients, display a drop in glucose metabolism in the brain [21, 22], prior to disease onset [22].

Glucose metabolism in the brain is coupled between glia and neurons [23]. According to the lactate shuttle hypothesis, most of glucose is taken up by glia and metabolised to lactate, which is then shuttled via transporters to neurons, where it is converted back to pyruvate and fed into the Tricarboxylic Acid Cycle (TCA) cycle for ATP production (16) (Fig 1A). An increase in lactate transport may therefore provide crucial fuel to meet the energy requirements of distressed neurons and allow them to better cope with the increase of toxic elements associated with disease.

**Fig 1.**
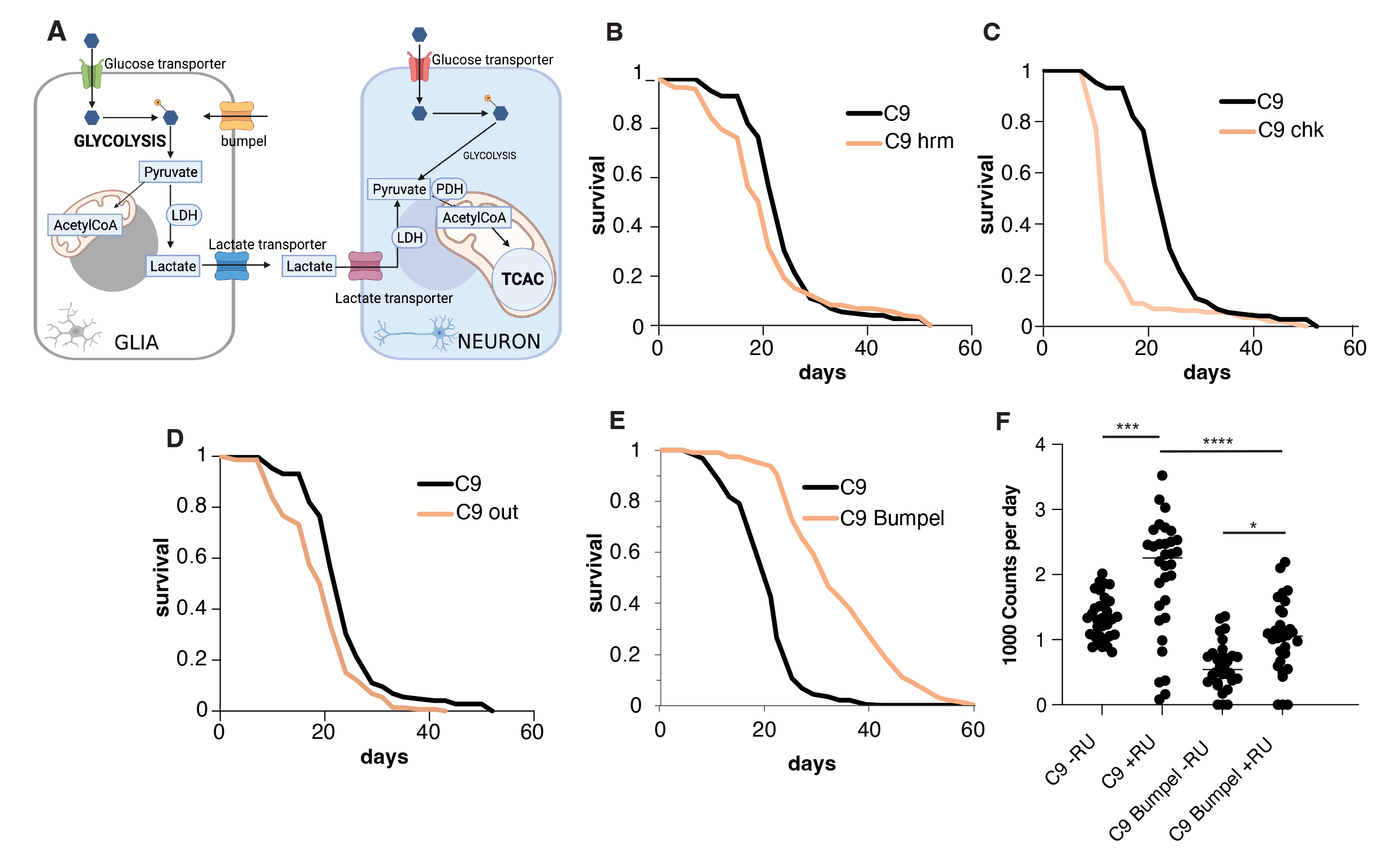
Bumpel over-expression in flies. (A) Diagram of the coupling of glucose metabolism between glia and neurons. Glia are mostly glycolytic, import glucose and break it down to pyruvate, which is exported out of glia, via transporters and into neurons, where it is converted back to pyruvate and then to Acetyl CoA for entry into the tricarboxylic acid cycle (TCA) cycle (Ldh=lactate dehydrogenase, Pdh=pyruvate dehydrogenase). (B) Survival curves of C9 expressing flies, induced with RU486, with (pink) and without (black) co-expression of *hrm*. p= 1.24E-56 by log rank test (C) Survival curves of C9 expressing flies, induced with RU486, with (pink) and without (black) co-expression of *chk.* p=4.9E-21 by log rank test (D) Survival curves of C9 expressing flies, induced with RU486, with (pink) and without (black) co-expression of *out* p=8.8E-6 by log rank test (E) Survival curves of C9 expressing flies, induced with RU486, with (pink) and without (black) co-expression of *bumpel*, p=1.75E-07 by log rank test (F) Activity counts of flies expressing C9 with and without bumpel (+RU) and their respective uniduced controls (-RU). p values of one-way ANOVA followed by Tukey’s multiple comparison test *=0.0199, ***=0.0002, ****< 0.0001 Please note the hrm, chk and out lifespans were run all at the same time and are displaying the same control. Genotypes: (A) *UAS-36 (G4C2), elavGS, UAS-36 (G4C2), elavGS/UAS-hrm,* (B) *UAS-36 (G4C2), elavGS, UAS-36 (G4C2), elavGS/UAS-chk,* (C) *UAS-36 (G4C2), elavGS, UAS-36 (G4C2), elavGS/UAS-out*, (D) *UAS-36 (G4C2), elavGS, UAS-36 (G4C2), elavGS/UAS-bumpel*.

Monocarboxylate metabolites, such as lactate and pyruvate, have been increasingly shown to play signaling roles too [24], suggesting they might play more broad ranging roles in the brain homeostasis, beyond their role in energy metabolism [24-26]. Lactate and pyruvate are shuttled across cell membranes by transporters, these can belong to two families: facilitated transporters (of the family of Solute Carrier 16 (SLC16) transporters some of them with proton co-transport) or active transporters (of the family of SLC5 sodium coupled transporters) [27].

The role of monocarboxylates in disease has not been tested in whole organisms, we decided to test whether increased monocarboxylate import could modulate neurodegenerative disease linked pathologies.

To address this question, we turned to flies. Glucose metabolism in *Drosophila*, similarly to higher eukaryotes, is also coupled between neurons and glia [28], with glia being mostly glycolytic and neurons carrying out mostly oxidative phosphorylation. We use our fly model of the C9orf72 repeat expansion (C9) [29], which was one of the first models of disease to report the toxicity associated with C9 repeats [29], with expression of 36 or 100 repeats leading to a dramatic shortening in lifespan and eye degeneration, due to the generation of toxic DPRs [29].

In this work, we over-expressed putative monocarboxylate transporters and found that, *bumpel,* a member of the conserved family of the SLC5A sodium coupled monocarboxylate transporters, can dramatically rescue the phenotype of C9 expressing flies, and lead to a reduction in the levels of specific DPRs. We show that *bumpel* is indeed a lactate and pyruvate transporter and that its overexpression is associated with the clearance of late stage autolysosomes, which accumulate in response to C9 epxression.

Importantly over-expression of *bumpel* can also rescue an AD fly model, suggesting that an increase in monocarboxylate transporters in neurons could provide a broad ranging therapeutic intervention.

## Results

### Bumpel, a monocarboxylate transporter, rescues C9 flies

In order to examine the relevance of monocarboxylate import to neurodegeneration in vivo, we over-expressed members of each family of transporters, facilitated and active, to determine if monocarboxylate import could have an effect on C9 toxicity in adult flies. Our C9 fly model expresses 36 C9 repeats pan-neuronally only in adult flies thanks to the *Gal4-Gene-switch* (GS) driver [30]. The gene-switch driver is induced when flies are fed the drug RU486, which is done after flies have eclosed, ensuring the expression of the repeats is limited to post-mitotic adult neurons, thus eliminating any confounding developmental effects. This is especially important in metabolic studies, as neurons are mostly glycolytic during embryonic stages and only switch to oxidative phosphorylation after birth [31]. Flies over-expressing 36 C9 repeats have a very short lifespan [29] and locomotion defects [32].

To test whether monocarboxylate import could ameliorate C9 associated phenotypes, we searched for transporters in the FlyORF collection of over-expression lines, which contains over 2000 lines, all inserted in the same site, as we wanted the lines to be comparably expressed. We identified lines over-expressing three genes homologous to the SLC16A family of transporters, *hermes* (*hrm), outsiders (out)* and *chaski* (*chk)*, and one SLC5 transporter, *bumpel*. We over-expressed all these in adult neurons of our fly model expressing 36 C9 repeats, to see if they could ameliorate the lifespan shortening phenotype (Fig 1B-E). Whereas over-expression of the homologous genes to the SLC16A family *hrm, out* and *chk* did not rescue C9 toxicity (Fig 1B, C and D), the over-expression of *bumpel* leads to a dramatic increase in lifespan from medians of 20 days, to nearly 30 days medians (Fig 1E). We confirmed the expression of *bumpel* by qPCR (Fig S1). Since both bumpel and C9 are driven by the same Gal4-GS driver, it is possible that binding to the UAS-bumpel promoter is titrating out the driver protein, leading to a drop in C9 expression and therefore, a phenotypic rescue. To confirm this is not the case we generated flies carrying an empty vector inserted in the same location as the FlyORF constructs used above, this also contains the binding sites for the driver and therefore could account for any dilution effects. Expression of the “empty” vector has no effect on lifespan of C9 expressing flies (Fig S1B), confirming that the rescue is due to bumpel and not to a non-specific dilution of the driver.

Bumpel was also able to rescue a 49 pure repeat fly model [33] (Fig S2), confirming its ability to rescue a variety of repeat lengths. Bumpel could also rescue motor phenotypes of C9 flies. C9 expressing flies display an early hyperactivity phenotype, with increased activity around day 4, this was brought back down by co-overexpression of *bumpel* (Fig 1F), showing that bumpel could rescue an early as well as late phenotypes associated with C9 expressing flies.

*Bumpel* is most homologous to human *SLC5A12* (solute carrier family 5 member 12) with 34% identity (56% similarity); *SLC5A5* (solute carrier family 5 member 5), with 34% identity (53% similarity); and *SLC5A8* (solute carrier family 5 member 8), with 35% identity (56% similarity). These proteins are known to promote transport of several substrates, including iodide (SLC5A5), pyruvate and lactate (SLC5A8 and SLC5A12).

*Bumpel*, together with its paralogues *kumpel* and *rumpel,* is expressed in glia in flies, where it is thought to promote transport of substrates across the brain [34].

### Bumpel reduces DPR accumulation

Bumpel’s rescue could be either due to a specific rescue of C9 toxicity or to a general improvement of organismal health. To distinguish between these possibilities we over-expressed *bumpel* in the neurons of a wild type fly and found that the lifespan was slightly shortened (Fig 2A), suggesting that the rescue is specific to C9 toxicity and is not due to a general improvement in health. C9 toxicity in flies is primarily driven by DPR accumulation [29], we therefore measured the levels of DPRs in C9 flies. We found that both GR and GP (Fig 2B and C) were reduced in C9 fly heads, indicating that the rescue is due to a downregulation of toxic DPR proteins. This was not due to a general drop in transcription from the elav promoter as the levels of the Gal4GS driver were actually increased by the presence of bumpel (Fig S3), or to general reduction in translation efficiency as levels of an mCD8-GFP protein, also driven by the elav-Gal4GS driver was not affected by bumpel expression (Fig 2D).

**Fig 2.**
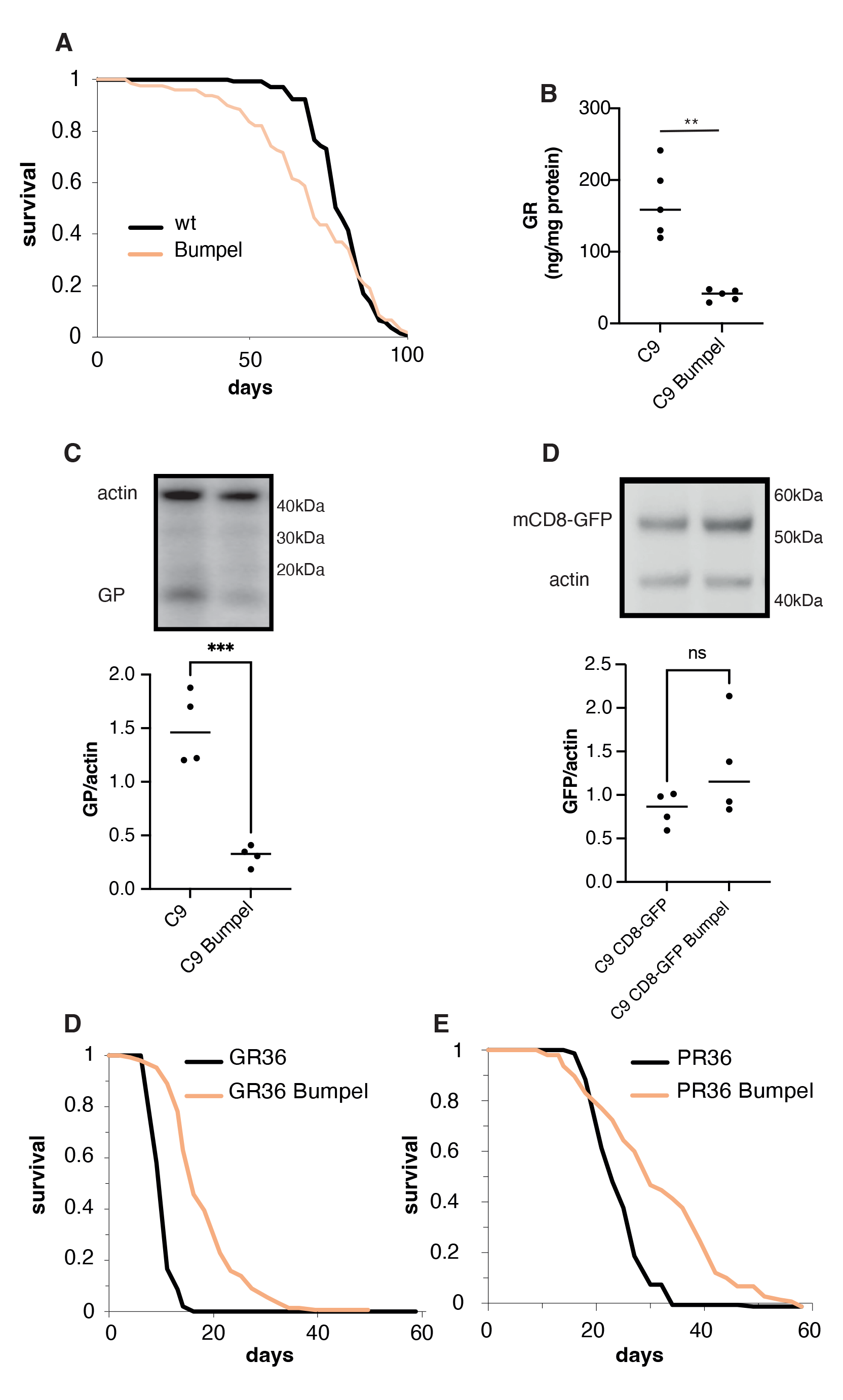
Over-expression of bumpel reduces DPR levels. (A) Survival curves of flies over-expressing *bumpel* (+RU) and their controls (-RU) during normal ageing. p=0.06 by log rank (B) GR levels in fly heads over-expressing C9 alone or with bumpel after 7 days of induction, measured by ELISA. p=0.0005 by unpaired t-test (C) GP levels in fly heads over-expressing C9 alone or with bumpel after 7 days of induction, measured by western blot. p=0.0005 by unpaired t-test (D) mCD8GFP levels in fly heads over-expressing C9+mCD8GFP or with bumpel after 7 days of induction, measured by western blot. p=0.1723 by unpaired t-test. (E) Survival curves of flies expressing 36(GR) repeats with (pink) and without (black) bumpel. p=1.5E-44 by log rank test (F) Survival curves of flies expressing 36(PR) repeats with (pink) and without (black) bumpel. p=2.1E-74 by log rank test (G) Survival curves of flies expressing 36(GA) repeats with (pink) and without (black) bumpel. p=3.2E-9 by log rank test. Genotypes: (A) *elavGS/UAS-bumpel* (B, C) *UAS-36 (G4C2), elavGS, UAS-36 (G4C2), elavGS/UAS-bumpel (D) UAS-36 (G4C2)/UAS-mCD8GFP, elavGS, UAS-36 (G4C2)/UAS-mCD8GFP, elavGS/UAS-bumpel* (E) *UAS-36GR, elavGS, UAS-36 GR, elavGS/UAS-bumpel* (F) *UAS-36PR, elavGS, UAS-36PR, elavGS/UAS-bumpel. (G) UAS36GA, elavGS, UAS36GA, elavGS/UAS-bumpel*

The decrease in DPR levels could either be due to bumpel specifically affecting RAN translation from the C9 repeats or due to an increase in DPR degradation. To distinguish between these possibilities, we checked the ability of bumpel to affect the lifespan of flies expressing and translating GR alone, from a standard ATG start codon, without any underlying repetitive sequence able to undergo RAN translation [29]. GR is the most toxic DPR produced by the C9 sense repeats. We found that bumpel was able to strongly rescue GR toxicity (Fig 2E). We also found that bumpel is able to rescue PR toxicity (Fig 2F), however bumpel over-expression was not able to rescue GA toxicity (Fig 2G), suggesting the rescue is limited to arginine containing DPR proteins.

### Bumpel is a pyruvate importer

Based on sequence homology, bumpel is predicted to transport both lactate and pyruvate [34], to confirm this we drove bumpel in larval fly salivary glands expressing lactate sensor Laconic (Fig 3A) or the pyruvate sensor Pyronic (Fig 3B), and exposed isolated tissue to either lactate or pyruvate. In both cases the salivary glands expressing bumpel showed a faster increase of monocarboxylate import (for lactate control vs bumpel: 0.1073 min^−1^ vs. 0.1717 min^−1^ ; for pyruvate: 0.0037 min^−1^ vs. 0.010 min^−1^) and to a higher extent (for lactate over-expression of bumpel a 74% increase and for pyruvate a 152% increase). This demonstrates that bumpel can transport both lactate and pyruvate.

**Fig 3.**
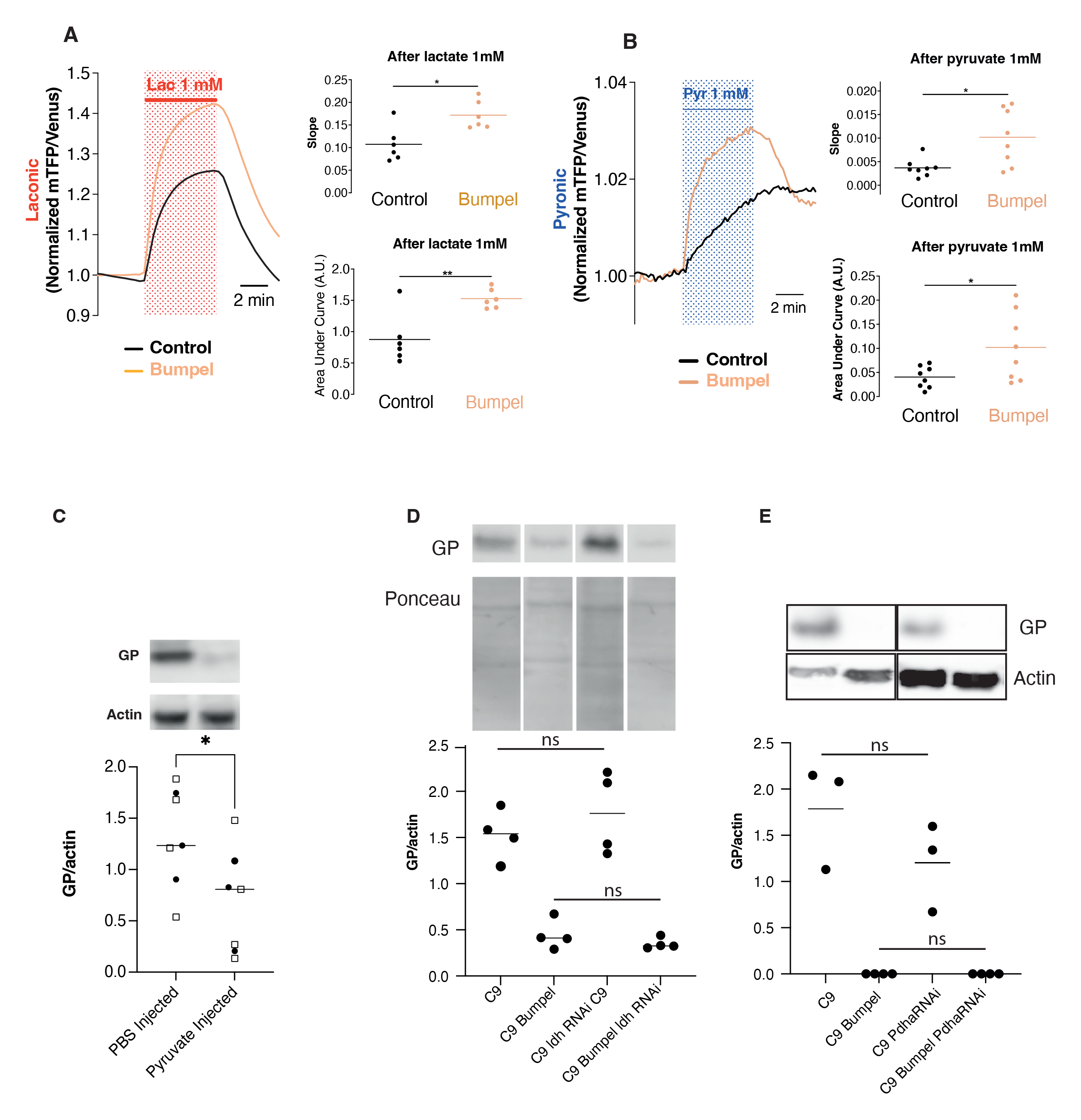
Bumpel leads to pyruvate import. (A) and (B) Lactate and pyruvate transport were measured using the lactate sensor Laconic (A) or the pyruvate sensor Pyronic (B). (A) Salivary glands from third instar larvae expressing Laconic and bumpel or its control were isolated and exposed to 1 mM lactate. Traces show the mean fluorescence ratio of mTFP/Venus obtained from 6 independent experiments (6 animals, total of 71 cells in control animals and 77 cells in animals over-expressing bumpel). Slope was calculated from the first 2 min of recording in the presence of lactate. p=0.026 by Mann Whitney test Area under the curve was calculated from the fluorescence obtained during the 5 min of lactate exposure. p=0.004 by unpaired t-test (B) Salivary glands expressing Pyronic and bumpel or its control were isolated and exposed to 1 mM pyruvate. Traces show the mean fluorescence ratio of mTFP/Venus obtained from 8 independent experiments (69 cells in control animals and 68 cells overexpressing bumpel). Slope were calculated from the first 2 min of recordings in the presence of pyruvate. p=0.026 by Mann Whitney test Area under the curve was calculated from the fluorescence obtained in 5 min of pyruvate exposure. p= 0.0346 by unpaired t-test. (C) Western blot of GP levels, relative to actin, in flies injected with 300 mM ethyl pyruvate and 2 M sodium pyruvate, or PBS as a control, and allowed to recover for 24 hours. Two different experimental replicates were combined in the statistical analysis (marked by different symbols), and then compared by t-test, *p= 0.0360 (D) GP levels in C9 and C9 bumpel fly heads with or without downregulation of Ldh by RNAi (E) GP levels in C9 and C9 bumpel fly heads with or without downregulation of Pdha by RNAi. GP levels in both D and E were compared by one way ANOVA followed by Šídák’s multiple comparisons test. All westerns samples shown for each panel were run on the same gel Gentoypes: (A) *UAS-36 (G4C2), actinGal4/UAS-Laconic* (B) *UAS-36 (G4C2), actinGal4/UAS-Pyronic (C) UAS-36 (G4C2), elavGS, (D) UAS-36 (G4C2), elavGS UAS-36 (G4C2)/LdhRNAi, elavGS, UAS-36 (G4C2), elavGS/UAS-bumpel, UAS-36 (G4C2)/LdhRNAi, elavGS/UAS-bumpel, (E) UAS-36 (G4C2), elavGS, UAS-36 (G4C2)/Pdha RNAi, elavGS, UAS-36 (G4C2), elavGS/UAS-bumpel, UAS-36 (G4C2)/PdhaRNAi, elavGS/UAS-bumpel*

To check whether an increase in pyruvate import was responsible for the drop in DPRs we injected a mixture of sodium pyruvate and ethyl pyruvate, which will cross membranes more readily and therefore the blood brain barrier, and reach the CNS better. Injection of adult 36R expressing flies induced on RU for 1 or 2 days, followed by 1 day recovery, showed a reduction in GP levels (Fig 3C). These experiments suggest that it is the increased import of pyruvate that leads to a drop in DPR levels. Pyruvate in the cell can be readily metabolized to Acetyl-CoA by Pyruvate dehydrogenase (Pdha) and enter the TCA cycle, it can also be converted to lactate by lactate dehydrogenate (ldh). We tested if imported pyruvate needed to be metabolized to exert its effect by knocking down ldh and Pdha by RNAi. Knock-down of both ldh and Pdha had no effect on the ability of bumpel to down-regulate GP levels (Fig 3D, E, S4), suggesting it is the metabolite itself that is leading to DPR degradation, not a down-stream product of its metabolisation.

### Bumpel modulates DPR levels via autophagy

An increase in intracellular pyruvate uptake has been recently shown to acidify the cytoplasm [35, 36], and induce mitophagy and autophagy [35-37], leading to protection against cell death in Parkinson’s disease cell culture models [35, 36].

Upregulation of mitophagy has been shown to lead to a drop in proteins associated with neurodegenerative disease which localise to mitochondria: experiments in worms have shown that pharmacological increase in mitophagy can indeed reduce the levels of Aß [38], which has been found to localise to mitochondria [38]. GR has also been shown to localise to mitochondria [39], therefore an upregulation in mitophagy could lead to the changes in toxic protein levels in C9.

We found that bumpel over-expression led to an increase in Pink and Parkin mRNAs, two key regulators of mitophagy of damaged mitochondria [40] (Fig 4A and B) and a drop in mitochondria copy number in heads, measured by the ratio of mtDNA to nuclear DNA (Fig 4C, S5A). This was confirmed by looking at levels of mito-GFP, which localises to mitochondria and found it to be reduced in the presence of bumpel, whereas a membrane localised GFP, mCD8-GFP was not affected (Fig 4D), suggesting a potential increase in mitophagy, or a drop in mitochondrial biogenesis in bumpel over-expressing flies.

**Fig 4.**
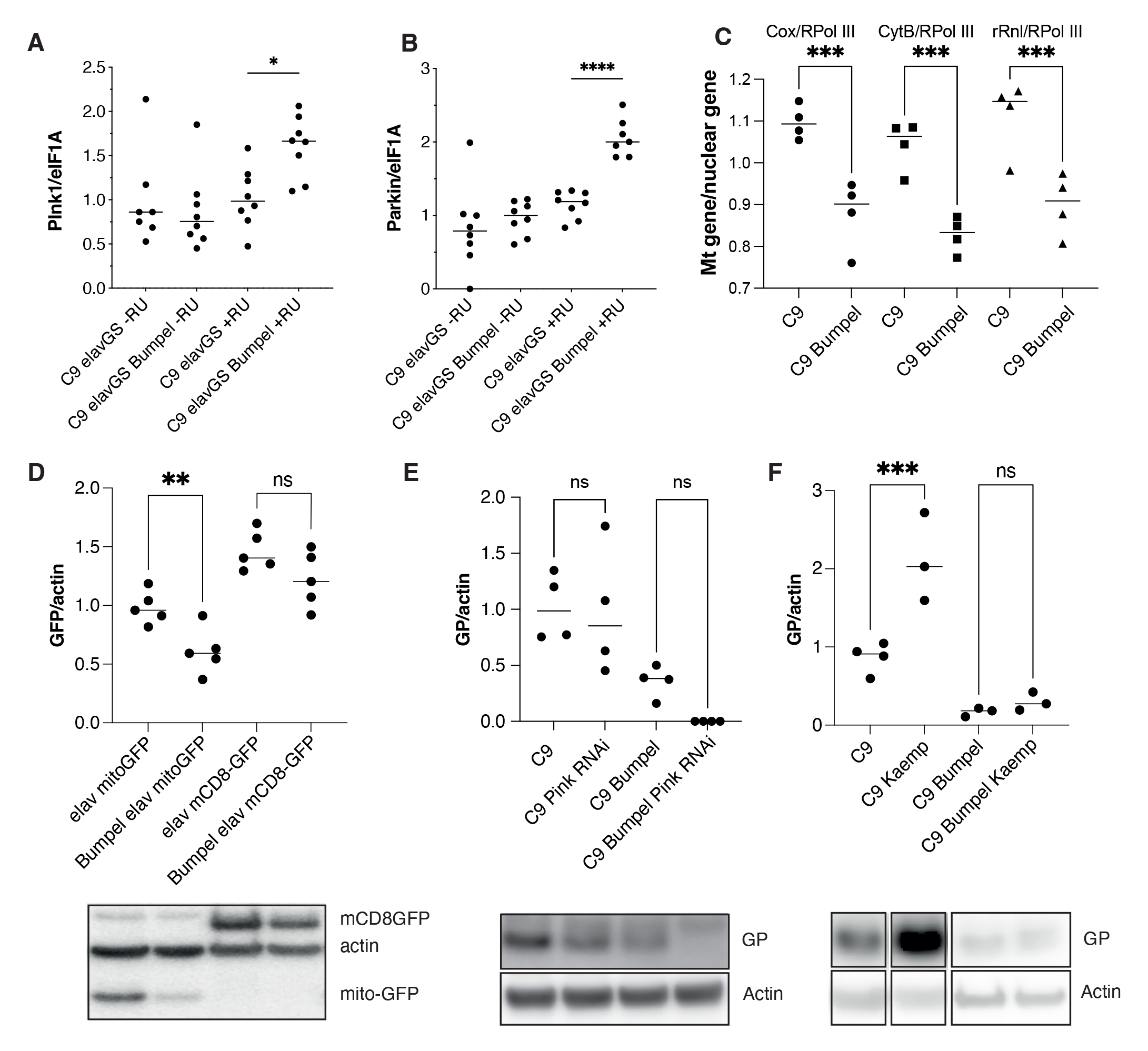
(A) Pink mRNA levels measured by qPCR, relative to eIF1A in C9 and C9 bumpel expressing flies (+RU) and controls (-RU). Genotype: *UAS-36 (G4C2), elavGS, UAS-36 (G4C2), elavGS/UAS-bumpel. *p<0.05 by* Šídák’s multiple comparisons test following one way ANOVA. (B) Parkin mRNA levels measured by qPCR, relative to eIF1A in C9 and C9 bumpel expressing flies (+RU) and controls (- RU). Genotype: *UAS-36 (G4C2), elavGS, UAS-36 (G4C2), elavGS/UAS-bumpel. ****p<0.0001 by* Šídák’s multiple comparisons test following one way ANOVA. (C) Mitochondrial copy-number measured by qPCR of mitochondrial DNA levels relative to nuclear DNA levels. ***p<0.0008 by Šídák’s multiple comparisons test following one way ANOVA. (D) GFP levels in fly heads expressing mitoGFP and mCD8GFP with and without bumpel **p= 0.0086 by student t-test (E) GP levels in C9 and C9 bumpel fly heads with or without downregulation of Pink1 by RNAi p=ns 0001 by Šídák’s multiple comparisons test following one way ANOVA (F) GP levels in C9 and C9 bumpel in heads of flies fed 50mg/L of kaempferol. ***p=0.0007 by Šídák’s multiple comparisons test following one way ANOVA. All westerns samples shown for each panel were run on the same gel Gentoypes: (A,B,C,E) *UAS-36 (G4C2), elavGS, (D) UAS-mitoGFP/elavGS, UAS-mCD8-GFP/elavGS, (E) UAS-36 (G4C2), elavGS, UAS-36 (G4C2)/Pink1 RNAi, elavGS, UAS-36 (G4C2), elavGS/UAS-bumpel, UAS-36 (G4C2)/Pink1RNAi, elavGS/UAS-bumpel*

We checked whether downregulation of Pink1, which regulates stress induced mitophagy [40], could affect DPR levels. Pink1 RNAi had no effect on GP levels in C9 or C9 bumpel expressing flies (Fig 4E), potentially suggesting that mitophagy does not play a role in regulating DPR levels. Pink1 is known to regulate stress induced mitophagy, however it’s role in basal mitophagy is less clear [41]. To confirm the role of mitophagy in modulation of DPR levels, we upregulated mitophagy pharmacologically with the potent inducer Kaempferol [42], this led to an increase in GP in 36R expressing flies, indicating that upregulation of mitophagy does not lead to a reduction of DPR levels (Fig 4F). This demonstrates that mitophagy upregulation is not a mechanism which can lead to a reduction in DPRs and therefore can not be responsible for the rescue observed in the bumpel expressing flies.

Import of pyruvate in fly salivary glands has also been shown to lead an increase in autophagy [43] and there are several lines of evidence from the literature to suggest that autophagy induction can indeed modulate DPR accumulation. C9orf72 protein has been shown to regulate endosomal trafficking and autophagy [44, 45] and a reduction in basal levels of autophagy has been observed in patient derived cell lines, due to C9orf72 haploinsufficiency [46] and this facilitates the accumulation of DPRs in cells [47]. Treating cells expressing 80 C9 repeats or individual DPRs with compounds inducing autophagy can lead to a reduction in DPRs [47, 48].

We decided to monitor autophagy in flies expressing C9 repeats. Autophagy is initiated by the engulfment of cellular material by autophagosomes, which, in later stages go on to fuse with lysosomes to form autolysosomes. We visualised autophagosomes with Atg8-mcherry and lysosomes with lysotracker dye. This allowed us to visualise both autophagosomes, only marked by Atg8-mcherry, and autolysosomes, marked by both Atg8-mcherry and lysotracker [49]. We found that expression of C9 repeats led to an increase in Atg8-mcherry vesicles, in particular autolysosomes, indicating a potential block in late stage autophagy (Fig 5 A, B and C), which could lead to accumulation of DPR proteins. Defects in late state autophagy and lysosomes have often been linked to neurodegenerative diseases [50], including in C9orf72 expansion models [51].

**Fig 5.**
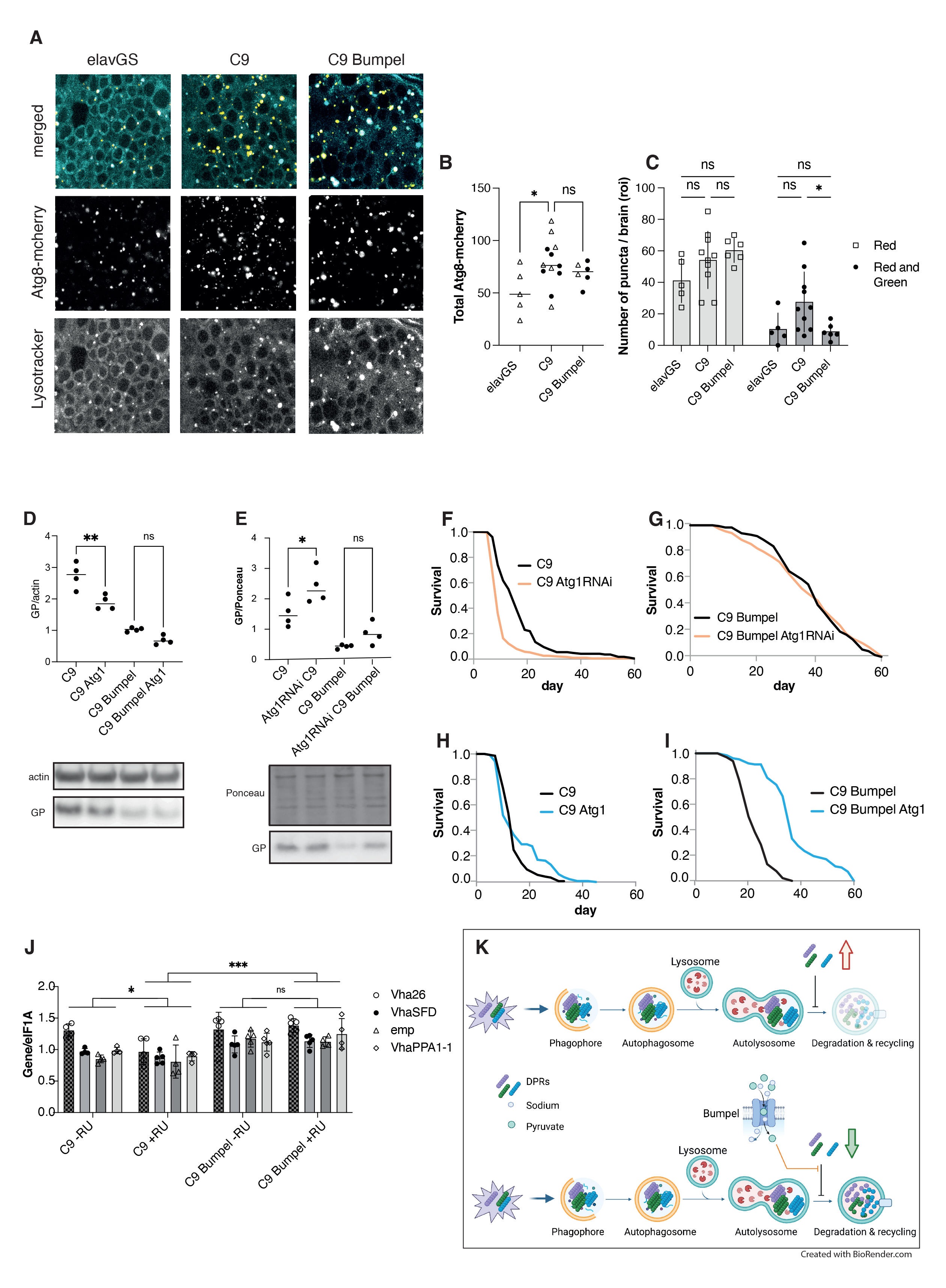
Autophagy can modulate GP levels. (A) Driver alone (elavGS) control, C9 expressing (C9) and C9+bumpel brains expressing Atg8-mcherry and stained with lysotracker, expression was induced with RU for 6 days (B) Total number of Atg8-mcherry vesicles, combined from 2 experiments, compared by t-test (C) Number of vesicles stained solely with Atg8-mcherry and those stained both with Atg8-mcherry and lysotracker, plotted as means with SEM error bars. Red only and red+green were compared separately by 1-Way ANOVA, *=0.042 by Šídák’s multiple comparisons test. (D) GP levels in fly heads over-expressing C9 with and without Atg1 and/or bumpel, measured by western blot. *p=0.0262 by Šídák’s multiple comparisons test following one way ANOVA (E) GP levels in fly heads over-expressing C9 with and without Atg1RNAi and/or Bumpel, measured by western blot. *p=0.023 by Šídák’s multiple comparisons test following one way ANOVA (F) Lifespan of C9 expressing flies with and without Atg1RNAi. p=1.43 E-19 by log-rank test (G) Lifespan of C9 bumpel expressing flies with and without Atg1RNAi. p=0.287 by log-rank test. (H) Lifespan of C9 expressing flies with and without Atg1. ADD LOG RANK (I) Lifespan of C9 bumpel expressing flies with and without Atg1. ADD LOG RANK (J) mRNA levels of Mitf targets in heads measured by qPCR relative to eIF1A. 2-way ANOVA followed by Tukey’s multiple comparison *p<0.05, ***p<0.0001. (K) Diagram of a model of the possible effect of DPRs and bumpel on autophagy. Autophagy initiates by the formation of a phagophore, engulfing aggregated proteins, including DPRs, to generate a autophagosome, which then fuses with a lysosome to generate an autolysosome, where the internal pH drops and proteases are induced, degrading organelles and proteins, and releasing the components back to the cytoplasm for re-cycling. DPR stall the maturation of the autolysosomes, components are not degraded and there is an increase in stalled autolysosomes, as well as DPRs, leading to a toxic feed-back loop (upper panel). In the presence of bumpel, pyruvate is imported, and this relieves the block in late autolysosomal maturation, allowing components to be degraded, autolysosomes to be cleared and leading to a drop in DPRs (lower panel). Genotypes (A, B, C and J) *UAS-mcherryAtg8, elavGS* ; *UAS-36 (G4C2)/UAS-mcherryAtg8, elavGS*; *UAS-36 (G4C2)/UAS-mcherryAtg8, elavGS/UAS-bumpel* (D, H and I) *UAS-36 (G4C2), elavGS, UAS-36 (G4C2), elavGS/UAS-bumpel; UAS-36 (G4C2), elavGS/UAS-Atg1; UAS-36 (G4C2), elavGS;UAS-bumpel/UAS-Atg1,* (E, Fand G) *UAS-36 (G4C2)/Atg1RNAi, elavGS, UAS-36 (G4C2)/Atg1RNAi, elavGS/UAS-bumpel*

We then checked if modulation of autophagy was able to alter the levels of DPRs. We over-expressed and downregulated the levels of Atg1, an essential mediator for autophagy initiation [52], in adult flies expressing 36 C9 repeats. We found that upregulation of Atg1 did indeed decrease the levels of GP in C9 expressing flies (Fig 5D), albeit not to the same extent as bumpel, whereas downregulation of Atg1 increases GP levels (Fig 5E). This suggests that autophagy can modulate DPR levels.

Modulation of Atg1 when bumpel was co-overexpressed, however, did not affect GP levels (Fig 5D, E), potentially suggesting that bumpel and Atg1 act in the same pathway and that bumpel modulates autophagy either downstream of Atg1 or independently of Atg1. QPCR confirmed that Atg1 was indeed reduced by the RNAi line used (Fig S6).

To confirm the role of autophagy in the modulation of C9 phenotypes, we down-regulated Atg1 in C9 flies and found that it shortened the lifespan of C9 expressing flies, demonstrating how autophagy can modulate C9 phenotypes (Fig 5F). In the presence of bumpel, Atg1 RNAi did not affect the lifespan of C9 expressing flies (Fig 5G), again suggesting bumpel and Atg1 act in the same pathway, and that bumpel is likely to activate autophagy either downstream or independently of Atg1. If bumpel was acting in the same pathway as Atg1, over-expression of both Atg1 and bumpel together could lead to a synergistic rescue. Atg1 over-expression in C9 flies alone did not affect phenotype (Fig 5H). However, when we over-expressed Atg1 in the presence of bumpel we found an increased rescue relative to bumpel alone, suggesting the two proteins are synergising together to rescue C9 phenotypes (Fig 5I).

We then monitored the ability of bumpel to modulate the autolysosome block observed in C9 expressing neurons. The bumepl transgene in the FlyORF stock is inserted into the ZH-86Fb site which carries a 3xP3-RFP construct, giving very strong red fluorescence in the brain. This RFP construct interferes with any imaging analysis. It is flanked by two loxP sites which we could excise using Cre recombinase. For image analysis we therefore used the bumpel stock no longer carrying the 3xP3-RFP.

We found that over-expression of bumpel did not decrease overall autophagosome numbers, but was able to specifically rescue the increase in autolysosomes (Fig 5A, C), suggesting it facilitates the clearance of autolysosomes, leading to degradation of DPRs.

The highly conserved transcription factor EB (TFEB) is a master regulator of lysosomal biogenesis and autophagy [53], and has been shown to facilitate the degradation of toxic proteins associated with neurodegenerative diseases in mammalian disease models [53], and in flies [54]. In C9 models nuclear-cytoplasmic transport defects lead to the mis-localisation of TFEB to the cytoplasm [51], leading to autolysosomal dysfunction. This would lead to a mis-regulation of TFEB target genes. The fly homologue of *TFEB*, *Mitf*, has been shown to co-ordinate the expression of a number of genes involved in lysosomal biogenesis, autophagy as well as lipid metabolism [54]. Of 38 genes described to be regulated by the Mitf, 16 are differentially expressed in a RNA seq dataset of fly heads expressing 100GR repeats [32], of these 16, 15 are downregulated (Table 1), suggesting that Mitf’s activity is downregulated in GR100 expressing flies. The toxicity in 36R expressing flies is mostly derived from the GR dipeptide repeat protein [29], and bumpel was able to rescue GR toxicity. We checked whether in C9 there was also a drop in Mitf targets and whether bumpel was able to restore the expression of Mitf target genes when co-overexpressed in C9 flies. Indeed, 4 Mitf targets, known to localise to the lysosome, were downregulated in response to C9 expression and their expression was increased in the presence of bumpel (Fig 5J), however a Mitf target not involved in lysosomal biogenesis, Cyp4d1, was not rescued (Fig S7) suggesting the rescue might be specific to Mitf targets involved in lysosomal biogenesis.

**Table 1.** Log fold change of TFEB targets genes in RNA seq of GR100 expressing flies vs control

| Gene | log2Fold Change | pvalue | padj |
| --- | --- | --- | --- |
| *Cyp4d1* | -0.86723 | 5.91E-05 | 0.000424 |
| *VhaAC39-1* | -0.210255 | 3.73E-06 | 3.32E-05 |
| *Vha36-1* | -0.184601 | 0.00071 | 0.004014 |
| *Vha26* | -0.182549 | 2.99E-06 | 2.71E-05 |
| *Vha55* | -0.175442 | 3.68E-05 | 0.000274 |
| *VhaSFD* | -0.174818 | 8.16E-05 | 0.00057 |
| *Vha100-2* | -0.170293 | 0.003694 | 0.017024 |
| *Vha13* | -0.164849 | 0.000585 | 0.003383 |
| *emp* | -0.15626 | 0.001682 | 0.008569 |
| *VhaM8.9* | -0.154655 | 0.000772 | 0.004328 |
| *Vha68-2* | -0.152605 | 0.002553 | 0.012427 |
| *VhaPPA1-1* | -0.144171 | 0.00521 | 0.022877 |
| *Vha44* | -0.143824 | 0.003112 | 0.014685 |
| *dor* | -0.138343 | 0.004453 | 0.020077 |
| *Vha16-1* | -0.129931 | 0.000381 | 0.002296 |
| *CtsB1* | 0.227224 | 1.19E-05 | 0.000097 |

These experiments suggest that bumpel over-expression leads to relief of a late lysosomal block within C9 expressing neurons, possibly via Mitf activation, leading to a drop in the levels of DPR proteins and amelioration of the C9 disease phenotype (Fig 5K).

### Bumpel rescues an Alzheimer’s disease fly model

We have found that an increase in pyruvate import is beneficial to an FTD/ALS fly model. Alzheimer’s disease is also characterised by a drop in glucose metabolism in patients’ brains [3] and late autolysosomal defects [55].

We wondered if increased transport of pyruvate could potentially be a broad-spectrum therapy and whether the over-expression of *bumpel* could be beneficial to an AD fly model. To explore this question, we over-expressed *bumpel* in a fly model expressing two copies of Aß42 [56], linked to signal peptide ensuring secretion.

These flies express high levels of Aß which can be secreted [57] and aggregate in Thioflavin S stained structures [56]. Similarly to our C9 model, we limit the expression of Aß to adult neurons, these flies have a shortened lifespan and a climbing defect as they age [58]. Interestingly, over-expression of *bumpel* ameliorated the lifespan and climbing phenotype of Aß expressing flies, and led to a reduction in Aß levels (Fig 6A, B and C), showing that bumpel can rescue an AD fly model too, and similarly to the C9 models, bumpel reduces toxic proteins associated with disease.

**Fig 6.**
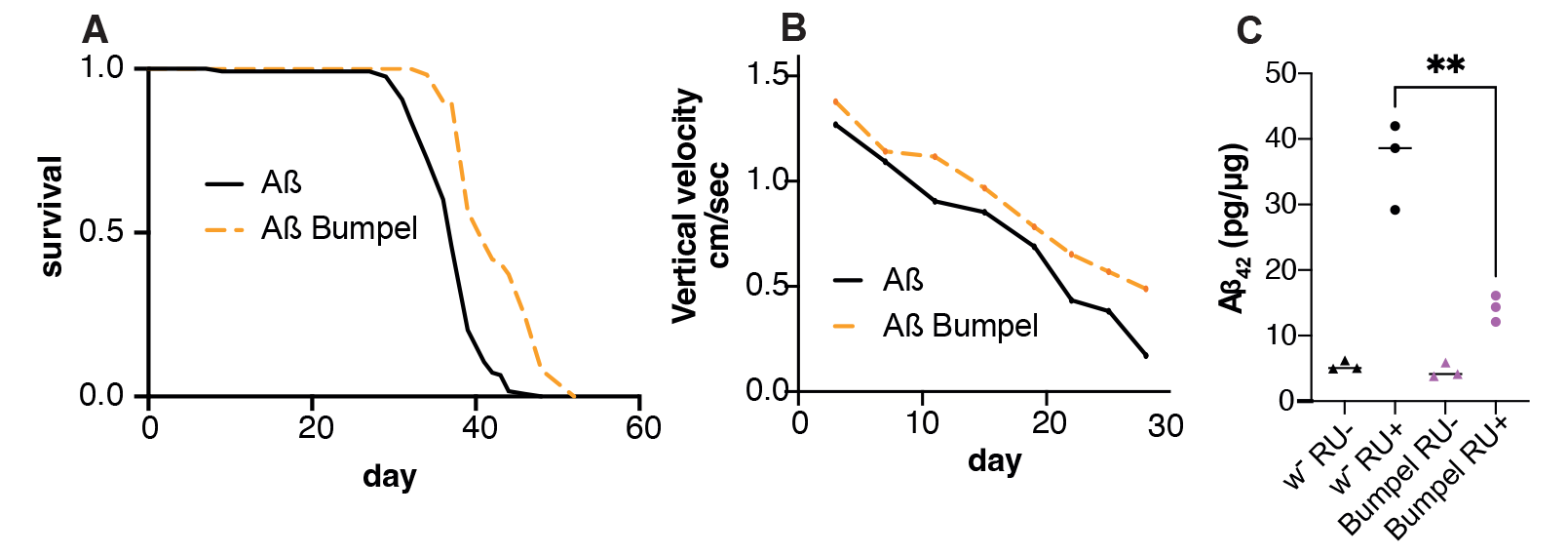
Bumpel over-expression rescues Aß expressing flies. (A) Survival curves of flies expressing Aß with (orange) and without (black) bumpel. p=1.3E-17 by log-rank test (B) Average climbing speed across the lifetime of flies expressing Aß with (pink) and without (black) bumpel. p=0.0016 by simple linear regression (C) Aß levels in fly heads over-expressing Aß alone or with bumpel after 7 days of induction, measured by ELISA. p=0.0033 by one way ANOVA followed by Dunnett’s multiple comparison test Genotypes: (A, B, C) *UAS-Aßx2, elavGS, and UAS-Aßx2, elavGS/UAS-bumpel*

## Discussion

Glucose hypometabolism of brain areas affected by disease is a common feature of many neurodegenerative diseases [59]. Glucose metabolism is tightly coupled between neurons and glia in brain, with lactate being shuttled from glia to neurons, to provide fuel for oxidative phosphorylation [23]. We have shown that upregulation of a monocarboxylate importer can ameliorate both C9orf72 repeat expansion toxicity and Aß toxicity in fly models. Both these diseases are caused by the accumulation of toxic peptides and in both cases the amelioration was linked to a drop in the toxic protein levels, suggesting that the phenotypic improvements are due to a downregulation of the toxic peptides.

The rescue was however not ubiquitous, toxicity due to the over-expression of poly(GA), one of the toxic DPRs associated with C9 repeats, was not rescued by bumpel over-expression, suggesting that the reduction of toxic peptides is specific to certain proteins.

Interestingly, over-expression of proton coupled lactate/pyruvate transporters did not rescue C9 phenotypes. It is possible that bumpel, since it is sodium coupled, would allow for constant import of lactate and pyruvate into neurons, even against a lactate/pyruvate concentration gradient, which proton coupled transporters might not allow. On the other hand, it is also possible that bumpel has a lower (or higher) affinity for substrates than other transporters, which could also explain the difference in behaviour. We show that the decrease in DPRs is associated with pyruvate import and that pyruvate itself is likely responsible for the rescue, and not a downstream metabolic process. Pyruvate has been shown to be able to modulate signalling pathways in the brain [26, 60], such as HIF1 transcription, or erythropoietin signalling pathway and plays a protective role following cerebral injury [61, 62]. Recently, import of pyruvate into cells has been associated with an induction of autophagy and mitophagy in cells [35-37] and flies [43]. We did indeed observe a reduction in mitochondrial copy number, when we co-overexpressed bumpel suggesting an increase in mitophagy. However, modulation of mitophagy did not appear to lead to a change in DPR levels, suggesting that it is unlikely to be a primary mechanism by which bumpel leads to DPR degradation.

Autophagic impairments have been associated with a number of AD models and an accumulation of autophagic vacuoles have been observed in patients’ brains [63]. Autophagic defects are thought to contribute to disease progression and facilitate the accumulation of amyloid beta [64], possibly because of faulty autolysosome acidification [55]. Several studies have shown that upregulation of autophagy can ameliorate Aß pathology and phenotypes in mice and cells [64-66], suggesting that an upregulation of autophagy can indeed lead to the clearance of Aß peptides. The same has been seen in C9 models, where defects in autophagy, especially linked to C9orf72 protein reduction in disease, can lead to an accumulation of DPR pathology [47], and upregulation of autophagy in cells can help with DPR clearance [47]. Increasingly, neurodegenerative disease have been linked to late autolysosomal defects [67] and these have also been observed in C9 models of disease [51].

We also observe an aberrant accumulation of autolysosomes in our C9 expressing fly neurons and over-expression of bumpel was able to promote autolysosomal clearance, suggesting it is acting late in autophagy to restore a healthy autophagic flux. This could be mediated by TFEB, a transcription factor regulating expression of many lysosomal genes [68], which has been shown to promote clearance of toxic peptide in a number of neurodegeneration models [69].

We confirmed C9 repeat expression impairs TFEB function, as seen in previous studies [51], and we find bumpel can restore the expression of fly TFEB target genes, specifically v-ATPase subunits, which could restore lysosomal function. How bumpel leads to a restoration of TFEB function will require further investigation.

Studies in human cells have linked expression of bumpel homologues and pyruvate import to histone deacetylase inhibition [70, 71]. TFEB’s activity is regulated by a number of acetylates and de-acetylases [72], so this could be a potential link, but it remains to be established if this is the case here.

We also find that increased autophagy, can reduce DPR levels, and that combined induction of early and late autophagy (by combining Atg1 with bumpel) leads to a synergistic rescue of C9 toxicity, suggesting that boosting autophagy with multiple interventions can be even more effective than a single intervention.

Pyruvate import might also be affecting neurons in other ways. Import of pyruvate into mitochondria could lead to an increase in oxidative phosphorylation and reactive oxygen species (ROS) formation, this, through a hormetic mechanism has been shown to boost cellular defence pathways [73]. An excessive increase in ROS formation could also explain the reduced lifespan when bumpel is over-expressed in a physiological ageing context, without disease.

Over-expression of bumpel is therefore a method to effectively increase autophagy within neurons affected by the toxic accumulation of neurodegenerative disease associated proteins, leading to their clearance. Bumpel in flies is predominantly a glial gene, which we are expressing in neurons, a tissue in which this gene is not normally expressed. Mis-expression of genes could present therapeutic challenges, however there a number of human homologues of bumpel, some, such as SLC5A8, are expressed in neurons, according to the human protein atlas, further studies are needed to pin-point which would be the best homologue to use in a human disease context.

This study overall points to a novel and unexplored role of pyruvate importers as a broad-spectrum modulator of toxic proteins associated with dementia, via neuronal autophagy enhancement, which could provide a novel therapeutic avenue.

## Materials and Methods

### Fly husbandry and stocks

All flies were reared at 25°C on a 12:12 light: dark cycle at constant humidity and on standard sugar-yeast medium (15g/L agar, 50 g/L sugar, 100 g/L autolysed yeast, 100g/L nipagin and 2ml/L propionic acid). Adult-onset, neuron–specific expression of UAS constructs was achieved as described in [29]. Briefly, 24-48 hours after eclosion female flies carrying a heterozygous copy of elavGS and at least one UAS-construct were fed SYA medium supplemented with 200µM mifepristone (RU486) to induce transgene expression.

Stocks used in this study:

UAS-36(G4C2), UAS-36GR, UAS-36PR, UAS-36GA have been described in [29], ElavGS was derived from the original elavGS 301.2 line [30] and obtained as a generous gift from Dr H. Tricoire (CNRS, France).

Atg1 GS10797 was obtained from the Kyoto Drosophila Genetic Resource Center. M{UAS-bumpel.ORF.3xHA.GW}ZH-86Fb, M{UAS-hrm.ORF.3xHA.GW}ZH-86Fb, M{UAS-out.ORF.3xHA.GW}ZH-86Fb, M{UAS-chk.ORF.3xHA.GW}ZH-86Fb were obtained from the flyORF collection [74]. W1118, Atg1 RNAi P{TRiP.HMS02750}attP40, UAS-mCD8::GFP, P{UAS-mito-HA-GFP.AP}2, Pdha RNAi P{TRiP.HMS06048}attP40/CyO, Pink1 RNAi P{TRiP.HMS01707}attP40, Ldh RNAi P{TRiP.HMS00039}attP2 were obtained from the Bloomington stock centre. The UAS-Aßx2 stock was a gift from Pedro Fernandez Funez (University of Minnesota) [56].

The ”empty” flyORF line was generated by injecting a pUAST plasmid in a w-; M{3xP3-RFP.attP}ZH-86Fb stock and selecting w+ flies.

To remove the 3xP3-RFP for image analysis, we crossed a UAS-bumpel, elavGS homozygous fly with a fly carrying Cre recombinase and a third chromosome balancer, and selected flies which had lost the RFP fluorescence in the eye, we confirmed these flies were still long lived.

### Lifespan analysis

All stocks used for lifespan analysis were back-crossed into a standard w1118 (for over-expression constructs) or v-w1118 stock (for Trip lines) for 6 generations to ensure homogenous genetic back-grounds. Male and female flies were allowed to mate and lay eggs for 24 hours on agar grape plates with yeast. The eggs were collected and seeded at standard density in 50ml bottles with SYA. After eclosion, flies were allowed to mate for 24-48 hours. At least 110-150 females of the appropriate genotype were split into groups of 15 and housed in vials containing SYA medium with or without drugs. Deaths were scored and flies tipped onto fresh food 3 times a week. Data are presented as cumulative survival curves, and survival rates were compared using log-rank tests. All lifespans were performed at 25°C.

### Analysis of activity and sleep

Individual, 4 day-old, mated, female flies were placed in 65 x 5 mm glass tubes containing standard 1xSYA, and activity was recorded using the DAM system (Drosophila Activity Monitoring System, TriKinetics, Waltham, MA) as described previously [75]. Flies were entrained to a 12:12 hour LD cycle at 25 °C and 65% humidity 24-36 hours before recording. Activity data are represented by mean values with their SEM. 32 flies were scored per genotype.

### Western Blotting

Protein samples were prepared by homogenising 10 fly heads per sample in 2x SDS Laemmli sample (4% SDS, 20% glycerol, 1 mM Tris-HCl pH6.8, 0 mM DTT with bromophenol blue) and boiled at 95°C for 5 minutes. Samples were separated on pre-cast 4-12% Invitrogen Bis-Tris gels (NP0322) and blotted onto PVDF in Tris-glycine buffer supplemented with 10% Methanol. Membranes were blocked in 5% milk in TBS-T (TBS with 0.05% Tween-20) for 1 hour at RT and then incubated with primary antibodies in TBS-T. Primary antibody dilutions used were anti-GP 1:3000 (a kind gift from Adrian Isaacs), anti-actin 1:10000 (Abcam ab1801), anti GFP (.Secondaries used were anti-rabbit and anti-mouse (Abcam ab6789 and ab6721) at 1:10000 dilutions for 1 hour at RT. Bands were visualized with Luminata Crescendo (Millipore) or Pierce ECL (Thermo Scientific) and imaged with ImageQuant LAS4000 (GE Healthcare Life Sciences). Quantification was carried out with ImageQuant software or ImageJ.

### qPCR

Total RNA was extracted from 10-15 fly heads per sample using Trizol (Invitrogen) and subsequently treated with DNAse I (Ambion) for DNA digestion. Equivalent amounts of RNA was then treated with TURBO DNase (Thermo Fisher Scientific). The RNA was then reverse transcribed using Superscript II (Invitrogen) with oligo(dT) primers. Quantitative gene expression analysis was performed on a QuantStudio 6 Flex Real-Time PCR System (Applied Biosystems) using SYBR-green technology (ABI). Relative quantities of transcripts were determined using the relative standard curve method normalized to eIF. Primer sequences can be found in table 1

**Table 1.**
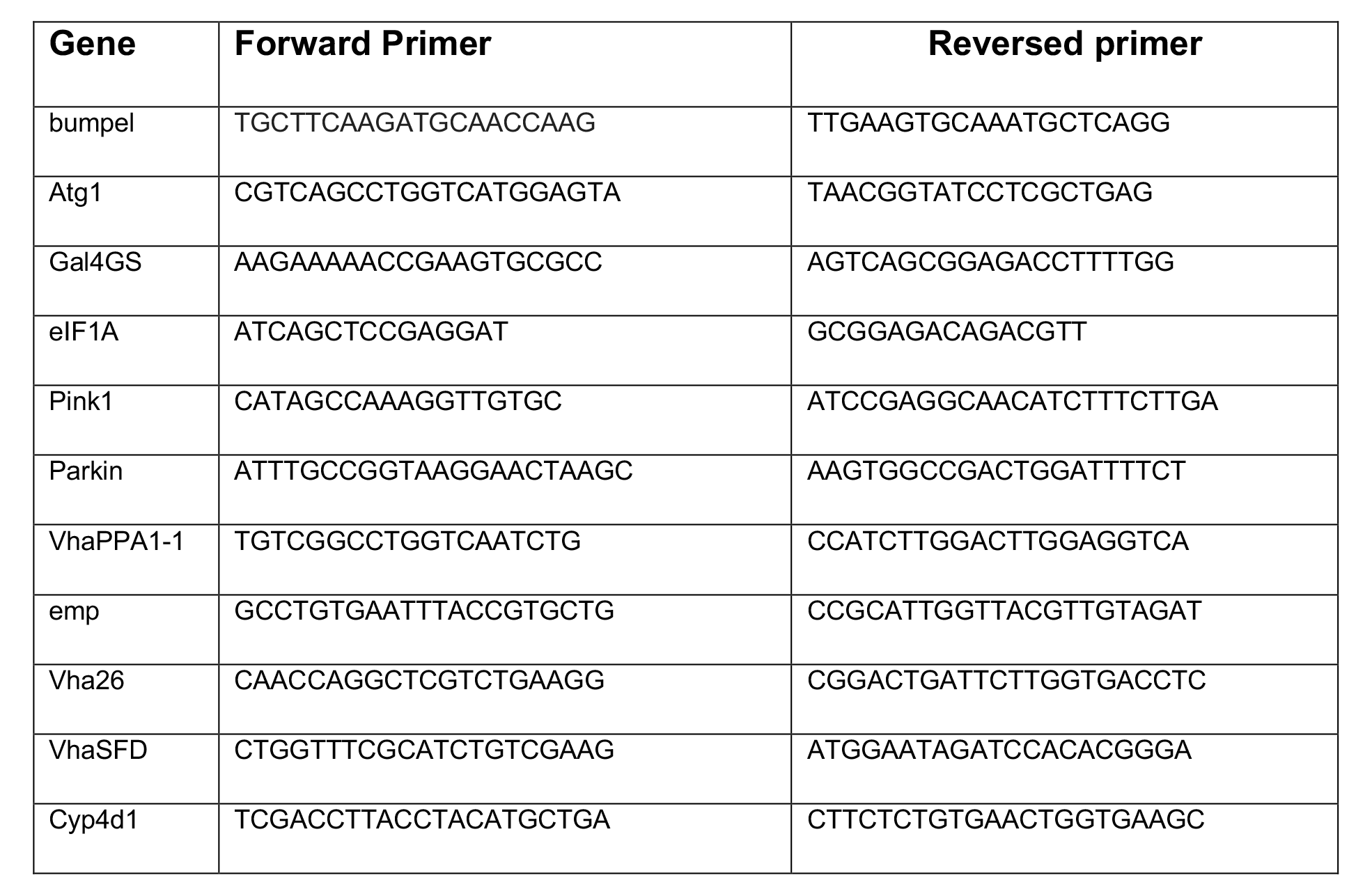
Primers for quantitative real-time PCR analysis

### Lactate and pyruvate measurements

Isolated salivary glands from control larvae (W1118) or over-expressing bumpel together with the FRET-based lactate sensor Laconic or the FRET-based pyruvate sensor Pyronic [76, 77] were placed in poly-L-lysine coated coverslips, mounted on a recording chamber (Live Cell Instruments, Seoul) and perfused with a solution composed by (in mM): 128 NaCl, 2 KCl, 1.5 CaCl2, 4 MgCl2, 5 trehalose, 5 HEPES and 35 sucrose (pH = 6.7) at 25°C. Changes in fluorescence signals were measured every 10 seconds before and after stimulation with 1 mM of lactate or pyruvate during 5 minutes in an Olympus BX61WI microscope equipped with DSU Spinning Disk Confocal System coupled to a DV2-emission splitting system (Photometrics, Tucson, Arizona) and a CCD camera Hamamatsu ORCA-R2 controlled by the Cell^R software (Olympus, Tokyo, Japan).

Images of Laconic and Pyronic fluorescence were obtained using an UMPlanFl X/0.5 water immersion objective, with an excitation of 438/24 nm and emissions at 488 nm (mTFP) and 540 nm (Venus) acquiring images every 10 seconds. In these images, we selected a region of interest in at least 8 cells of each salivary gland using ImageJ (1.52e, National Institutes of Health, USA), measuring the mean gray value minus the background. The result obtained was the fluorescence ratio between mTFP/Venus normalized to the mean value of 2 min before lactate or pyruvate exposure. Differences between control larvae and over-expressing Bumpel were assessed by the Mann-Whitney Sum Rank test (pairs) or Student’s t-test.

Significance was established with p-values less than 0.05.

### GR ELISA

10 female frozen heads per tube were homogenised in 2% SDS buffer (Cat No. 42- 801-8 0ML, Millipore Sigma) containing 1X RIPA buffer (Cat No. R0278, Sigma-Aldrich) and complete mini EDTA-free protease inhibitor cocktail (Cat No. 11836170001, Roche) at Room Temperature for about 30 sec or until the heads were no longer intact. Samples were then heated at 95C for 10 min. After centrifuged at 14000 rpm for 20 min at RT, the supernatants were collected in the new tubes and stored at −80˚C. The protein concentration was determined using Pierce BCA Protein Assay Kit (Cat No. 23325, ThermoFisher) according to the manufacturer’s manual.

Samples were diluted to the same concentration with homogenisation buffer and 25µL were loaded in duplicate in the 96-well Meso Scale Discovery (MSD) immunoassay plate. The singleplex MSD immunoassay was performed to measured poly-GR expression levels as described here [78].

### Fruit Fly Injection

3 day old C9 flies were induced with RU food for 24/36 hours. 37 nano-liters of PBS, 2 M NaCl (Sigma Aldrich) in PBS (control), and a mixture of 300 mM ethyl pyruvate (Sigma Aldrich) and 2 M sodium pyruvate (Sigma Aldrich) in PBS (pyruvate group) were then injected into the abdomen of each fly using Drummond “NANOJECT II” Automatic Nanoliter Injector (3-000-206A). All the flies were put back onto RU food for another 24 hours after injection and then flash-frozen by liquid nitrogen. Flies were processed for GP western as described above.

### Microscopy

Fly brains expressing Atg8-mcherry were dissected in Schneinder’s media and stained for 3 minutes with lysotracker (LysoTracker Red DND-99; Thermo Fisher Scientific; 1:2000), rinsed in Schneinder’s media and then directly visualised with a Zeiss Airyscan 880. Single confocal planes were taken for all images, and all settings were kept the same within an experiment. Images were analysed in ImageJ using the watershed plug-in. Objects were identified in the red channel and then the mean fluorescence of both the green and red channel was measured with the Analyse particle function. We exported the values to Excel, and particles with values of 0 for green fluorescence, were considered red-only, particles with values for each channel were considered as co-localising.

### Aß ELISA

Flies kept on RU486- or RU486+ food for 15 days were instantly frozen in liquid nitrogen. 10-15 fly heads were homogenised in guanidine extraction buffer (5M Guanidinium-HClSigma G3272, 50 mM Hepes Sigma H3375, 5 mM EDTA, protease inhibitor cocktail Roche Complete Mini). The plate was coated with first antibody solution (Clone 6E10, Covance SIG-39320, prepared 1:400 in 50 mM carbonate), incubated at 4°C overnight, washed, and blocked with 1:4 Block Ace solution in PBS (stock solution: 4 g of Block Ace powder in 100 mL ddH2O) for 2 hours. 2 µL of samples was added to each well containing 98 µL of diluent buffer (0.5 mL Triton-X, 0.5 mL 30 mg/mL PMSF EtOH solution, PBS 49 mL, protease inhibitors). Serial dilutions of Aβ 42 peptide (Meso Scale Discovery C01LB-2) were prepared in diluent buffer with extra guanidine extraction buffer (49 diluent: 1 guanidine extraction) and added to wells. The plate was incubated at 4°C overnight, washed and incubated with detection antibody solution (Clone 12F4, Covance SIG-39144, diluted 1:1000 in 1:10 Block Ace solution) for 2 hours. Then the plate was washed and incubated with 1:10,000 Neutravidin-HRP in PBS (Pierce 31001, stock 1mg/mL) for 1 hour. The plate was washed, and 100 µL of KPL TMB Microwell Peroxidase Substrate was added to all the wells. After 15 mins, 100 µL of 1M HCl was added to each well to stop reaction. The plate was read at 450 nm.

### Climbing assay

Mated female flies were split into vials containing RU486+ food (15 flies per vial, 5 vials per genotype), and were tipped onto new food three times a week. Twice a week, flies were tapped down to the bottom and allowed to climb. The whole process was recorded by a camera, and the videos were analysed by the automatic quantification system FreeClimber [79]. Vertical velocities of the fly populations were obtained and plotted in GraphPad Prism 9.

### Statistical analysis

Lifespans were compared by log-rank test, performed in Excel. Two way comparisons were performed by unpaired t-test and multiple comparisons, following a one way ANOVA, were performed either by Tukey, Šídák’s or Dunnett’s multiple comparisons test as appropriate. Comparisons between groups were performed by 2-Way ANOVA (Fig 5J). Tests were performed in GraphPad Prism 9.

When multiple replicates of the same experiment were combined, we compared the effect of the experimental replicate and the effect of the treatment by 2 way ANOVA. since the variability between experimental replicate was not significant, we combined the values from multiple experimental replicates together (this was done for Fig 5B and 5C).

## Acknowledgments

This work was supported by the ARUK Senior Fellowship ARUK-SRF2018A-003 (TN), ARUK-PG2016A-6 (AMI), MRC grant MR/V003585/1 (TN), the Alzheimer’s Society Studentship (550(AS-PhD-19b-015)), the European Research Council (ERC) under the European Union’s Horizon 2020 research and innovation programme (648716 – C9ND) (AMI) and the UK Dementia Research Institute (AMI), which receives its funding from UK DRI Ltd, funded by the UK Medical Research Council, Alzheimer’s Society and Alzheimer’s Research UK. FONDECYT-Iniciación 11200477 (to AGG) and FONDECYT Regular 1210586 (to JS).

We thank Nazif Alic for useful feedback during the preparation of the manuscript.

## SUPPLEMENTAL MATERIAL

**Supplemental Fig 1.**
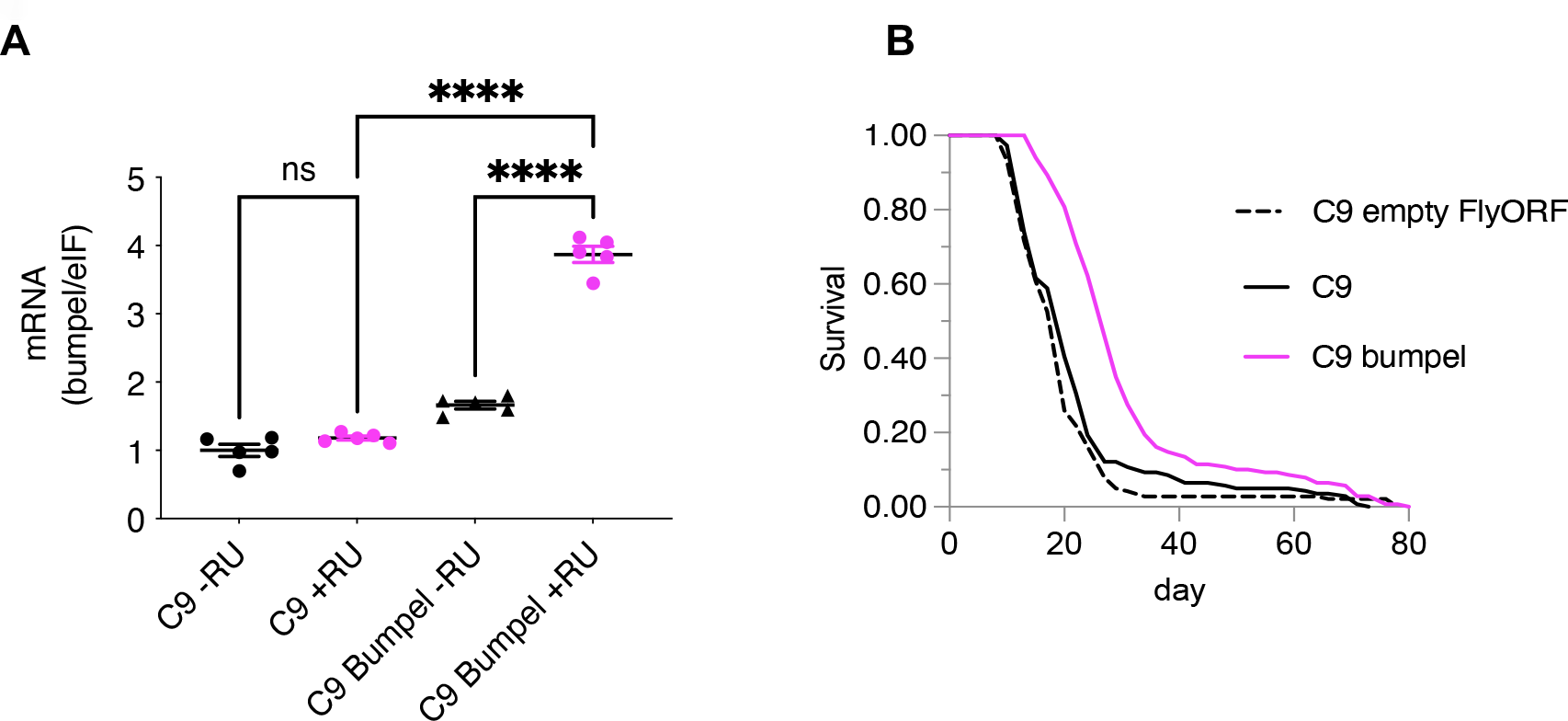
**A.** Bumpel mRNA levels measured by qPCR, relative to eIF in C9 and C9 bumpel expressing flies (+RU) and controls (-RU). Genotype: *UAS-36 (G4C2), elavGS, UAS-36 (G4C2), elavGS/UAS-bumpel. ****p<0.0001 by* Šídák’s multiple comparisons test following one way ANOVA **B.** Lifespan of 36R expressing flies with and without bumpel co-expression. Genotype: *UAS-36 (G4C2), elavGS; UAS-36 (G4C2), elavGS/UAS-empty FlyORF; UAS-36 (G4C2), elavGS/UAS-bumpel* p=3.4E-17 for comparison of C9 bumpel to C9 empty flyORF and p=1.3E-10 for C9 bumpel to C9.

**Supplemental Fig 2.**
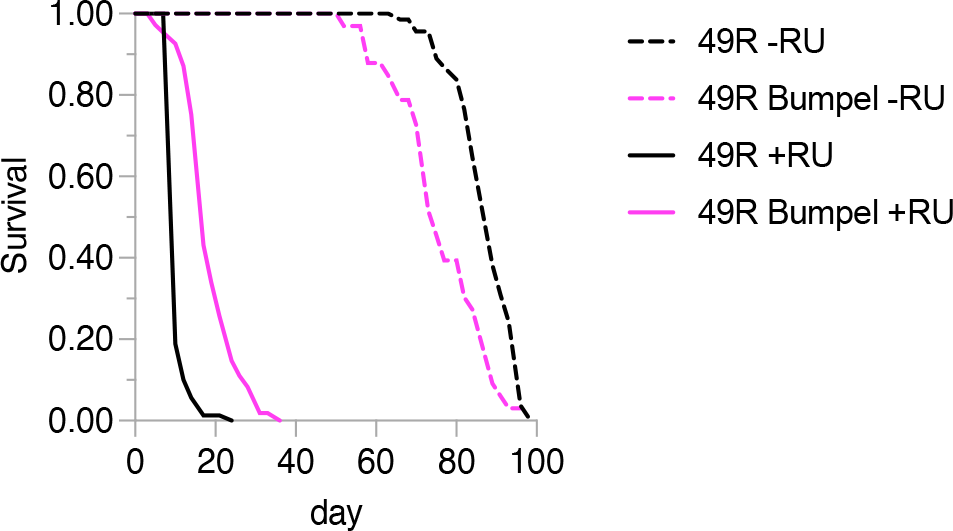
Lifespan of 49R expressing flies (+RU) with and without bumpel co-expression and their uninduced controls (-RU). Genotype: *UAS-49 (G4C2), elavGS, UAS-49 (G4C2), elavGS/UAS-bumpel* p=6.9E-36 for +RU comparisons.

**Supplemental Fig 3.**
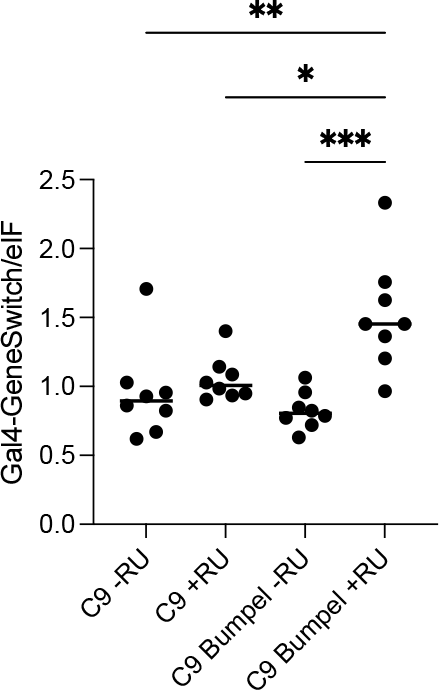
Gal4GS mRNA levels measured by qPCR, relative to eIF in control (- RU) and C9 and C9 bumpel expressing flies (+RU). ***p= 0.0002, **p= 0.0022, *p= 0.0142 *by* Šídák’s multiple comparisons test following one way ANOVA. Genotypes; *UAS-36 (G4C2), elavGS, UAS-36 (G4C2), elavGS/UAS-bumpel*

**Supplemental Fig 4.**
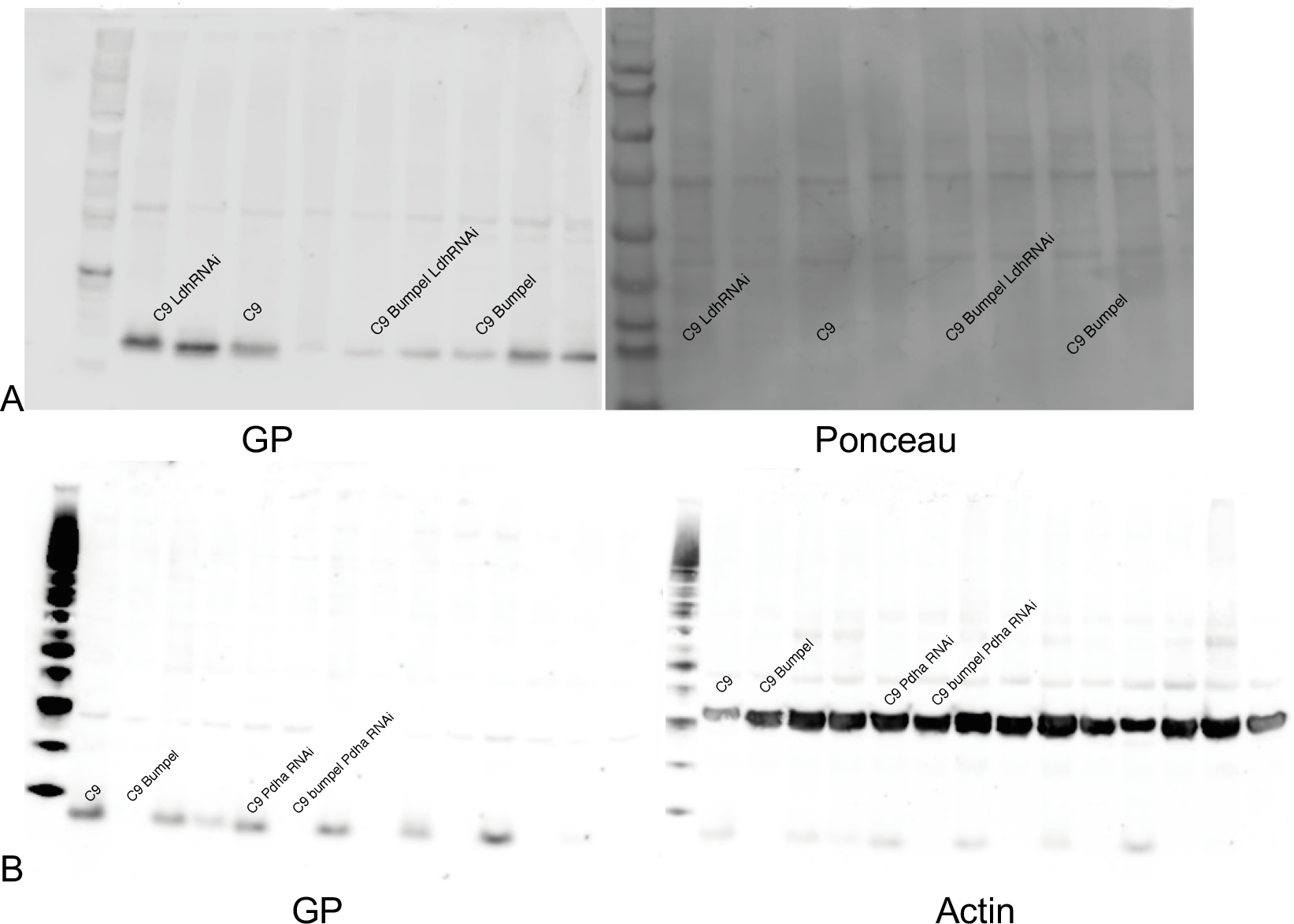
Full western blots for figures 3D (A) and 3E (B), with samples labelled.

**Supplemental Fig 5.**
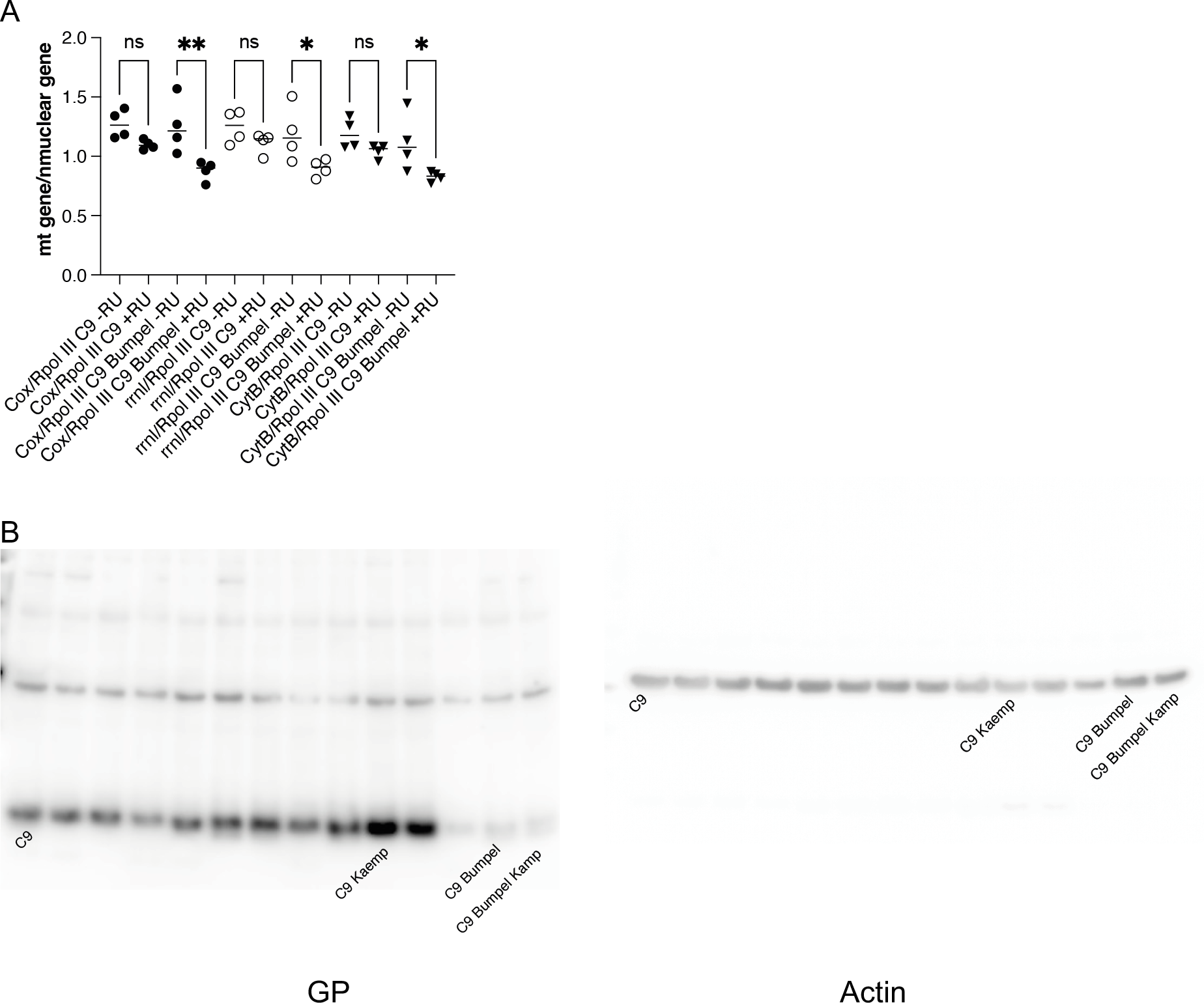
**A.** Mitochondrial copy-number measured by qPCR of mitochondrial DNA levels relative to nuclear DNA levels. *p<0.05, **p<0.005 by Šídák’s multiple comparisons test following one way ANOVA. +RU samples also shown in Fig 4C **B.** Images of the whole blots used in Fig 4F. Genotypes; *UAS-36 (G4C2), elavGS, UAS-36 (G4C2), elavGS/UAS-bumpel*

**Supplemental Fig 6.**
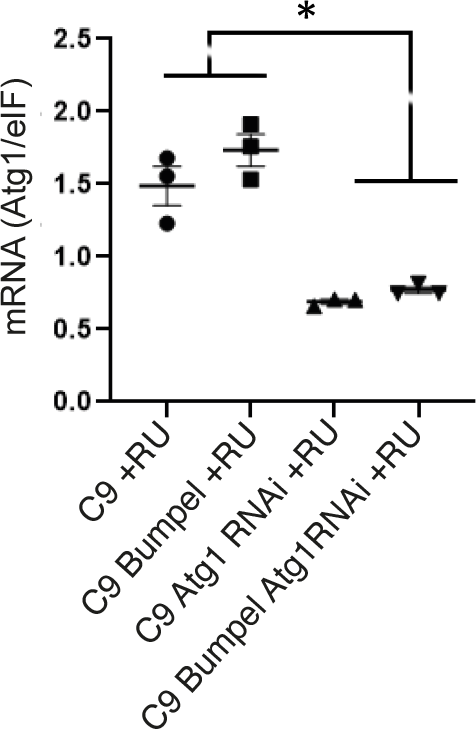
Atg1 mRNA levels measured by qPCR, relative to eIF in C9 Atg1RNAi and C9 bumpel Atg1RNAi expressing flies (+RU). *p<0.05 by by Šídák’s multiple comparisons test following one way ANOVA. Genotypes; *UAS-36 (G4C2), elavGS, UAS-36 (G4C2), elavGS/UAS-bumpel,UAS-36 (G4C2)/Atg1RNAi, elavGS, UAS-36 (G4C2)/Atg1RNAi, elavGS/UAS-bumpel,*

**Supplemental Fig 7.**
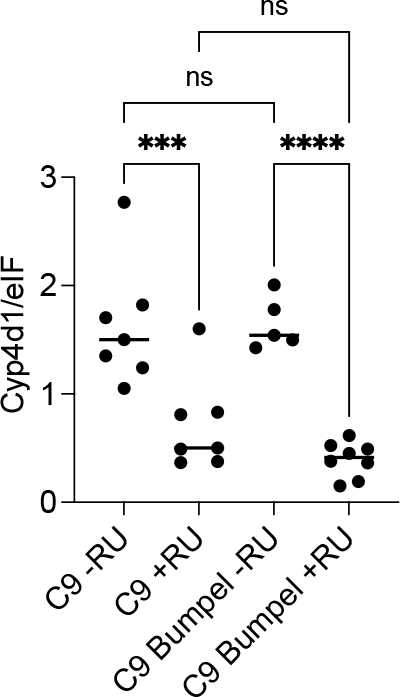
Cyp4d1 mRNA levels measured by qPCR, relative to eIF in control (- RU) and C9 and C9 bumpel expressing flies (+RU). ***p=0.001, ****p<0.0001 by by Šídák’s multiple comparisons test following one way ANOVA. Genotypes; *UAS-36 (G4C2), elavGS, UAS-36 (G4C2), elavGS/UAS-bumpel*

